# Stronger positive and purifying selection on X-linked genes in a species with paternal genome elimination

**DOI:** 10.64898/2026.09.14.751372

**Authors:** Sam Ebdon, Hollie Marshall, Christina Hodson, Clément Schneider, Laura Ross, Kamil S. Jaron

## Abstract

The faster-X effect predicts that X chromosomes may evolve more rapidly than autosomes due to selection on expressed recessive alleles. Quantifying the effect empirically usually requires numerous assumptions due to the impact of diplodiploid transmission dynamics, and so the relative contribution of dominance effects to sex chromosome evolution remains unclear. Springtails present a novel opportunity to ameliorate this. Male globular springtails have hemizygous X chromosomes, diploid autosomes, and eliminate their paternal genome during spermatogenesis. This leads to the autosomes matching the X chromosomes haplodiploid transmission. Hence, theoretically the X chromosomes and autosomes only differ in the fixation probability of male-beneficial alleles. We generated whole-genome resequencing and RNA-Seq data for the globular springtail *Allacma fusca*, to quantify the faster-X effect, leveraging this natural experiment. We find that after accounting for the deleterious distribution of fitness effects, and despite surprisingly depleted neutral X-linked diversity, the sex chromosomes show signals of both stronger purifying and positive selection, supporting the faster-X effect. Furthermore, we provide the first direct evidence for the loss of DNA methylation in Collembola. These results, obtained from a system where many of the confounding sex-autosome differences that typically obscure faster-X patterns are absent, support hemizygosity as a genuine driver of sex chromosome evolution and suggest that sex-linked genes could contribute disproportionately to adaptive potential.

## Introduction

Sex chromosomes evolve rapidly within populations and often contribute disproportionately to genetic differentiation between species (Coyne, 1992, Coyne & Orr, 1989, Haldane, 1922, Turelli & Orr, 1995). However, identifying the processes driving differences in divergence rates between sex chromosomes and autosomes remains challenging due to the large number of confounding factors involved (Charlesworth *et al*., 2018, Klein *et al*., 2021). In many diploid systems, sex chromosomes are hemizygous in one sex and diploid in the other. As a consequence of this, when new beneficial variants are at least partially recessive, they are more likely to fix on sex chromosomes than autosomes as they are expressed in the heterogametic sex, a phenomenon dubbed the faster-X (or Z) effect (Betancourt *et al*., 2002, Charlesworth *et al*., 1987, Vicoso & Charlesworth, 2006). Conversely, nearly neutral or weakly deleterious mutations are more likely to fix on autosomes, as hemizygosity increases the efficacy of purifying selection on sex-linked recessive variation for the same reasons (Vicoso & Charlesworth, 2006). Thus, assuming different chromosomes experience mutations with equivalent fitness and dominance effects, hemizygous sex chromosomes may experience both elevated rates of adaptive divergence and reduced rates of deleterious divergence relative to autosomes. Empirical support for the faster-X effect, however, is mixed across taxa (Ávila *et al*., 2015, Avila *et al*., 2014, Bechsgaard *et al*., 2019, Charlesworth *et al*., 2018, Dean *et al*., 2015, Kousathanas *et al*., 2014, Mank *et al*., 2010b, Meisel & Connallon, 2013, Mongue & Baird, 2024, Torgerson & Singh, 2006, Vicoso & Charlesworth, 2006), and it can be difficult to interpret where studies do not distinguish between adaptive and nonadaptive evolution due to their opposing effects on divergence (Charlesworth *et al*., 2018). Compounding this, empirical estimates of the dominance coefficients of new mutations are rare, having been obtained in only a small number of taxa including *Drosophila*, yeast, nematodes, and one plant and one human study (Agrawal & Whitlock, 2011, Di & Lohmueller, 2024, Huber *et al*., 2018, Kyriazis & Lohmueller, 2024, Phadnis & Fry, 2005). These estimates are typically confounded by the same factors that complicate faster-X inference.

This lack of consensus reflects the fact that in diplodiploid systems the dominance-based faster-X model is an oversimplification, as numerous other processes may alter substitution rates on sex chromosomes relative to autosomes (Charlesworth *et al*., 2018, Klein *et al*., 2021). Hemizygosity not only exposes recessive sex-linked alleles to selection but also reduces the expected effective population size of sex chromosomes to three-quarters that of autosomes, increasing the relative contribution of genetic drift (Vicoso & Charlesworth, 2006). This relative Ne expectation is frequently violated by variation in demography, mating system, or reproductive success, all of which can alter the dominance conditions expected to drive faster-X evolution (Mank *et al*., 2010b, Vicoso & Charlesworth, 2009). Mutation rates may also differ between chromosomes due to sex-biased transmission and asymmetry in germline mutation rates (Vicoso & Charlesworth, 2006), although this can be partially controlled for by normalising by neutral substitution rates (Vicoso & Charlesworth, 2009). In addition, recombination rates often differ between sex chromosomes and autosomes. In species with recombination in both sexes 2/3 of copies of the sex chromosomes recombine versus 4/4 copies of the autosomes. In species with an achiasmatic heterogametic sex 2/3 copies of the sex chromosomes recombine but only 2/4 copies of autosomes recombine. Thus, recombination is relatively reduced on sex chromosomes in species with recombination in both sexes (2/3 *<* 4/4) but can be elevated in systems with achiasmy in the heterogrametic sex (2/3 *>* 2/4), altering the effective span of selective sweeps and background selection (Charlesworth *et al*., 2018). Further asymmetries arise from the haplodiploid transmission of the X chromosome, which forces alleles to alternate between sexes across generations (Klein *et al*., 2021), and biases the reproductive value of X-linked loci towards females (Charlesworth *et al*., 1987). Finally, sex chromosomes and autosomes may differ in gene content (Meisel & Connallon, 2013, Rice, 1984, Vicoso & Charlesworth, 2006), suggesting that the distribution of fitness and dominance effects of new mutations may also differ between chromosomes. Together these factors complicate attempts to derive general expectations for differences in evolutionary rates between sex chromosomes and autosomes.

One way to reduce the complexity introduced by the numerous assumptions required when quantifying the faster-X effect is to take advantage of natural experiments provided by alternative genetic systems (Normark, 2003, Ross *et al*., 2022). Species with paternal genome elimination (PGE, also known as pseudoarrhenotoky) are particularly informative (Burt & Trivers, 2006, Herbette & Ross, 2023, Normark, 2003). Elimination occurs at various stages in different species, but some in particular, such as gall midges (Bantock, 1970, White, 1946), fungus gnats (Du Bois, 1933, Gerbi, 1986, Goday & Esteban, 2001, Haig, 1993), and globular springtails (Dallai *et al*., 2001, 2000, Jaron *et al*., 2022), X chromosomes are eliminated early in embryogenesis but autosome elimination is delayed until spermatogenesis (Dallai *et al*., 2000). The autosomes are expected to maintain biparental expression, but as males do not transmit the chromosomes inherited from their fathers, the autosomes are effectively haploid with respect to inheritance (see Jaron *et al*., 2022, Ross *et al*., 2022). This changes the transmission dynamics of autosomes from a diplodiploid to a haplodiploid pattern of inheritance, and therefore they match the X chromosome(s) (Normark, 2003) (Fig. 1). Under this mode of PGE, effective population sizes and recombination rates are equivalent between all chromosomes, male and female contributions to each generation are balanced, and X-linked alleles alternate between the sexes at the same frequency as autosomal alleles, yet males retain haploid X chromosomes (Klein *et al*., 2021). Thus, this system allows us to test whether the expression of X-linked (partially) recessive alleles leads to faster evolution relative to autosomes (Table 1), solely due to selective effects (Hitchcock *et al*., 2024, Orr, 2001).

**Figure 1:**
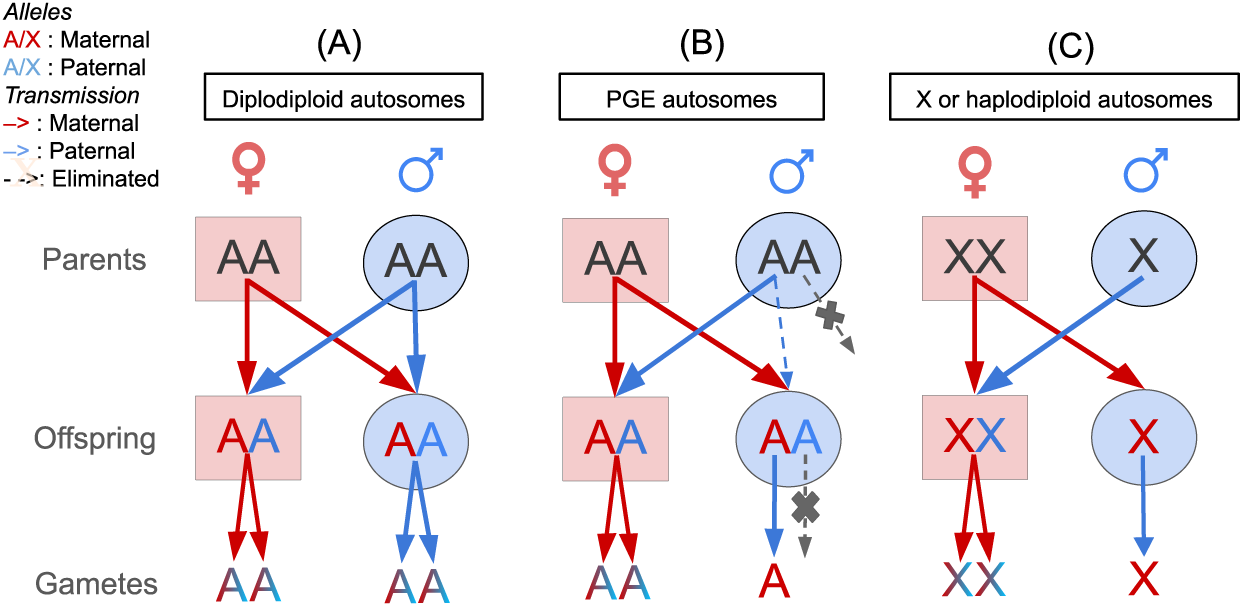
A) In diplodiploids, autosomes inherited from either parent can be transmitted to offspring of either sex by either parent. B) As in diplodiploids, under paternal genome elimination (PGE) females pass on either their maternal or paternal autosomes to both daughters and sons. In contrast, males eliminate their paternally inherited chromosomes, and only pass on their maternally inherited autosomes to their offspring. As a result, autosome transmission each generation matches the haplodiploid pattern of X chromosomes, but all adults retain diplodiploid autosomes in the soma. C) X chromosome transmission in females is as in diplodiploids, but males must pass on their only copy to daughters. This schematic is a simplification as the paternal X does get passed on to males but it is eliminated early in development. Females are shown in red squares and males are shown in blue circles. Red and blue letters represent chromosomes inherited from the individuals mother and father respectively, while red and blue lines represent transmission from mothers and fathers respectively to offspring. Dashed lines show lineages destined for elimination and are crossed at elimination. Lines originate between chromosomes where either chromosome can be transmitted or at specific chromosomes when transmission is constrained.

**Table 1:** Comparison of evolutionary features expected to drive X-autosome differences in diplodiploid systems and those with paternal genome elimination (PGE). Ticks indicate features contributing X-autosome differences. Crosses indicate no X-autosome differences.

| Feature | PGE | Diploid |
| --- | --- | --- |
| Haploid expression | ✓ | ✓ |
| Gene-specific selection | ✓ | ✓ |
| Effective population size | ✗ | ✓ |
| Recombination rates | ✗ | ✓ |
| Sex copy-number bias | ✗ | ✓ |
| Sex alternation bias | ✗ | ✓ |

To date, few studies have leveraged species with ‘atypical’ transmission dynamics to quantify variation in chromosome evolution while minimising confounding factors. Baird *et al*. (2025) estimated evolutionary rates in Sciarid flies, a family of fungus gnats that undergo PGE. They showed that the X chromosome diverges more slowly than the autosomes, likely as a consequence of more effective purifying selection reducing the weakly deleterious substitution rate. Jaquiéry *et al*. (2018) utilised a similar natural experiment in aphids, which produce males asexually by eliminating one X chromosome copy in the germline. They observed evidence for faster-X effects via relaxed purifying selection on silenced genes in combination with weak positive selection. Like PGE, this mechanism equilibrates transmission dynamics between X chromosomes and autosomes, though here the X is shifted to match the autosomes rather than vice-versa.

We investigate the selected and neutral components of diversity and divergence to quantify rates of X chromosome evolution in the globular springtail *Allacma fusca* using another member of Sminthurinae subfamily, *Sminthurus viridis*, as an outgroup. Springtails are small and abundant non-insect arthropods found in a wide range of habitats including soil, leaf litter, trees, fresh water, and rockpools (Hopkins, 1997). Globular springtails of the order Symphypleona exhibit paternal genome elimination (Dallai *et al*., 2001, 2000, Jaron *et al*., 2022) and have two X chromosomes (X_1_0X_2_0) (Dallai *et al*., 2004, 2000) which take up a significant portion of their genome (*≈* 28% or 103 Mb in *A. fusca*). Importantly, globular springtails eliminate their X chromosomes early in embryogenesis, likely before the onset of embryonic gene expression, such that paternally-derived X-linked genes are unlikely to ever be expressed. This makes them an ideal PGE system for investigating the faster-X effect. However, this assumes that the paternal autosomes are expressed in males, which we will seek to confirm given that some other PGE species silence the paternal subgenome (de la Filia *et al*., 2021, Herbette & Ross, 2023, Ross *et al*., 2022).

We use a chromosome level assembly for *Allacma fusca* (GCA 947179485.1, Jaron *et al*. (2023)) and *Sminthurus viridis* (GCA 965194885.1), 12 wholegenome sequencing (WGS) (Jaron *et al*., 2022) and generated 20 RNA-Seq datasets to quantify the site-frequency spectra at non-synonymous and synonymous sites within genes and for intergenic regions. Both species are symphypleonans in the same family — Sminthurinae — and share the same PGE system. We confirm that male globular springtails express both parental haplotypes, as predicted by their mode of paternal genome evolution. This corroborates cytogenetic observations that while paternal chromosomes are silenced in certain PGE clades, they are not heterochromatinized in *Allacma* (Dallai *et al*., 2000, Herbette & Ross, 2023). We then ask the following questions. Firstly, we test whether PGE in globular springtails is modulated by the epigenetic mechanism, DNA methylation. Secondly, we ask whether genetic diversity is lower on X chromosomes due to increased purifying selection. Thirdly, we infer deleterious distributions of fitness effects (dDFE) and quantify adaptive rates of evolution using *DFE-alpha* (Eyre-Walker & Keightley, 2009, Keightley & Eyre-Walker, 2007) to investigate whether purifying and positive selection is stronger on X-linked genes than autosomal genes and hence estimate the efficacy of haploid selection. Finally, we conduct a differential expression analysis between sexes to annotate sex-biased genes and to investigate the potential for more effective adaptive and purifying selection in male-biased genes.

## Results

We annotated 24,243 genes (18,542 autosomal and 5,701 X-linked) in *A. fusca* using a combination of RNA-Seq reads and known arthropod proteins and 41,460 genes (34,637 autosomal and 6,823 X-linked) in *S. viridis* using *A. fusca* proteins and the arthropoda database only using BRAKER (Gabriel *et al*., 2024). After annotating transposable elements using the EarlGray pipeline (Baril *et al*., 2024), we estimated approximately 70% and 64% of the genome is intergenic in *A. fusca* and *S. viridis* respectively and both were approximately 35% repetitive. Interestingly while gene density was relatively uniform across chromosomes in *S. viridis* (31% to 37%), *A. fusca*’s two largest autosomes are relatively gene poor (17% and 22% vs 32% - 37% across remaining chromosomes). In addition, *A. fusca*’s largest autosome is syntenic with *S. viridis*’s second smallest, suggesting non-genic expansions in *A. fusca* (Fig. S1). Repetitive regions and transposable elements were excluded from further analyses.

### Male globular springtails express both parental autosomal haplotypes despite PGE

We quantified the expression of annotated genes using RSEM (Li & Dewey, 2011) with RNA-Seq reads obtained from whole pooled and individual adult springtails. A PCA of gene expression (PC1 94%, PC23%) separated the sexes on PC1 revealing one of the pooled RNA-Seq datasets contain a mixture of males and females (Fig. S2). After excluding the mixed-sex library, differential expression analyses were conducted using 9 male (four pools of three and five individuals) and all 10 female (five pools of three and five individuals) RNA-Seq libraries. Log2 fold changes (log2FC) were estimated using DESeq2 (Love *et al*., 2014), with apeglm shrinkage applied prior to classification of sex-biased genes. We considered genes as sex-biased if they showed a log2FC*>*1 in expression towards either sex and a FDR adjusted p-value below 0.05. Of all genes tested, 5,642 were male-biased and 2,243 female-biased, with male-biased genes enriched on the X chromosomes (Fisher’s exact test, OR = 0.589, p *<* 0.0001) (Fig. S3). Interestingly, the composition of genes with sex-biased expression differed between the two X chromosomes (Two-sided X-squared = 56.834, df = 2, p-value *<* 0.0001). Female-biased genes were enriched on X1 (20%, OX359249.1) relative to X2 (14%, OX359250.1), while the opposite was true for male-biased genes (32% and 40%), and unbiased proportions were similar (48% and 46%). We investigated dosage balance and compensation using TMM-normalised TPMs and comparing male-female and X-autosome ratios (Fig. 2A). We filtered for genes expressed in both sexes and excluded genes outside the 5% and 95% quantiles in each sex. This resulted in 6,774 autosomal and 2,784 X linked genes. Males and females are approximately balanced on both the autosomes and the sex chromosomes with a slight skew towards males (Autosomes: Mann-Whitney U, median male = 2.93, median female = 2.71, median M:F = 1.17, p *<* 0.0001; X: Mann-Whitney U, median male = 3.11, median female = 2.84, median M:F = 1.18, p *<* 0.0001), likely driven by very strongly expressed male-biased genes. When comparing sex chromosomes separately, genes on the X1 showed near-complete dosage balance between the sexes (median M:F = 0.997), whereas genes on the X2 exhibited a male-biased shift in expression (median M:F = 1.22), suggesting the male-biased skew is driven by X2. Unusually, we see slight (*≈* +5%) hyperexpression of both X chromosomes in both males and females (Male: Mann-Whitney U, mean A:X = 0.945, p *<* 0.0001; Female: Mann-Whitney U, mean A:X = 0.954, p *<* 0.0001), which has also been seen in another PGE system (Baird *et al*., 2025). Excluding sex-biased genes reduced but did not eliminate X hyperexpression (Mann-Whitney U; M A:X = 0.942, p *<* 0.0001; F A:X 0.989, p *<* 0.0001).

**Figure 2:**
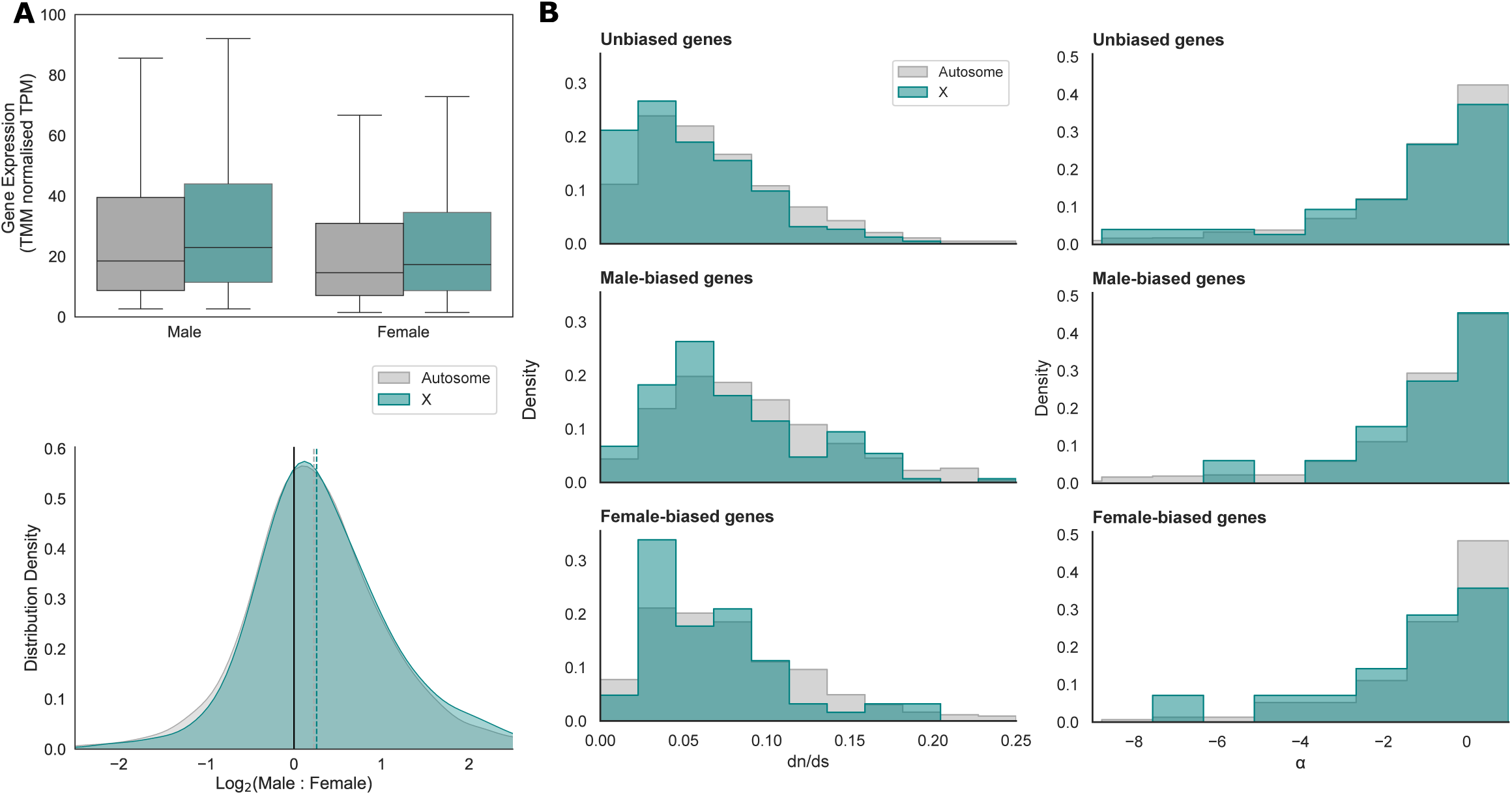
A) Dosage compensation (upper) and dosage balance (lower) on the X chromosomes (teal) and autosomes (grey) in male-biased and female-biased genes in the globular springtail *Allacma fusca*. While the sexes show approximate dosage balance, both males and females show hyperexpression of X-linked genes relative to autosomal genes. B) KDE plots of *dn/ds* (left) and positive selection strength, *α* (right, estimated using the McDonald-Kreitman test) using single-copy orthologs between *Allacma fusca* and *Sminthurus viridis*. Plots are shown for unbiased, male-biased, and female-biased genes respectively from top to bottom on the X chromosomes (teal) and autosomes (grey).

We tested if male globular springtails express both parental haplotypes on the autosomes as expected given their PGE dynamics by quantifying the relative coverages of reference and alternative alleles at heterozygous sites called in five single-sample male and female RNA-Seq datasets. Relative coverages on male autosomes are intermediate and match both female autosomes and X chromosomes (Male A: 0.449 [0.324, 0.574], Female A: 0.0453 [0.328,0.578], Female X: 0.452 [0.326, 0.578]) demonstrating that males are expressing paternally inherited autosomes (Fig. 3). The average number of X-linked heterozygous SNPs called in males was approximately 26% that of females.

**Figure 3:**
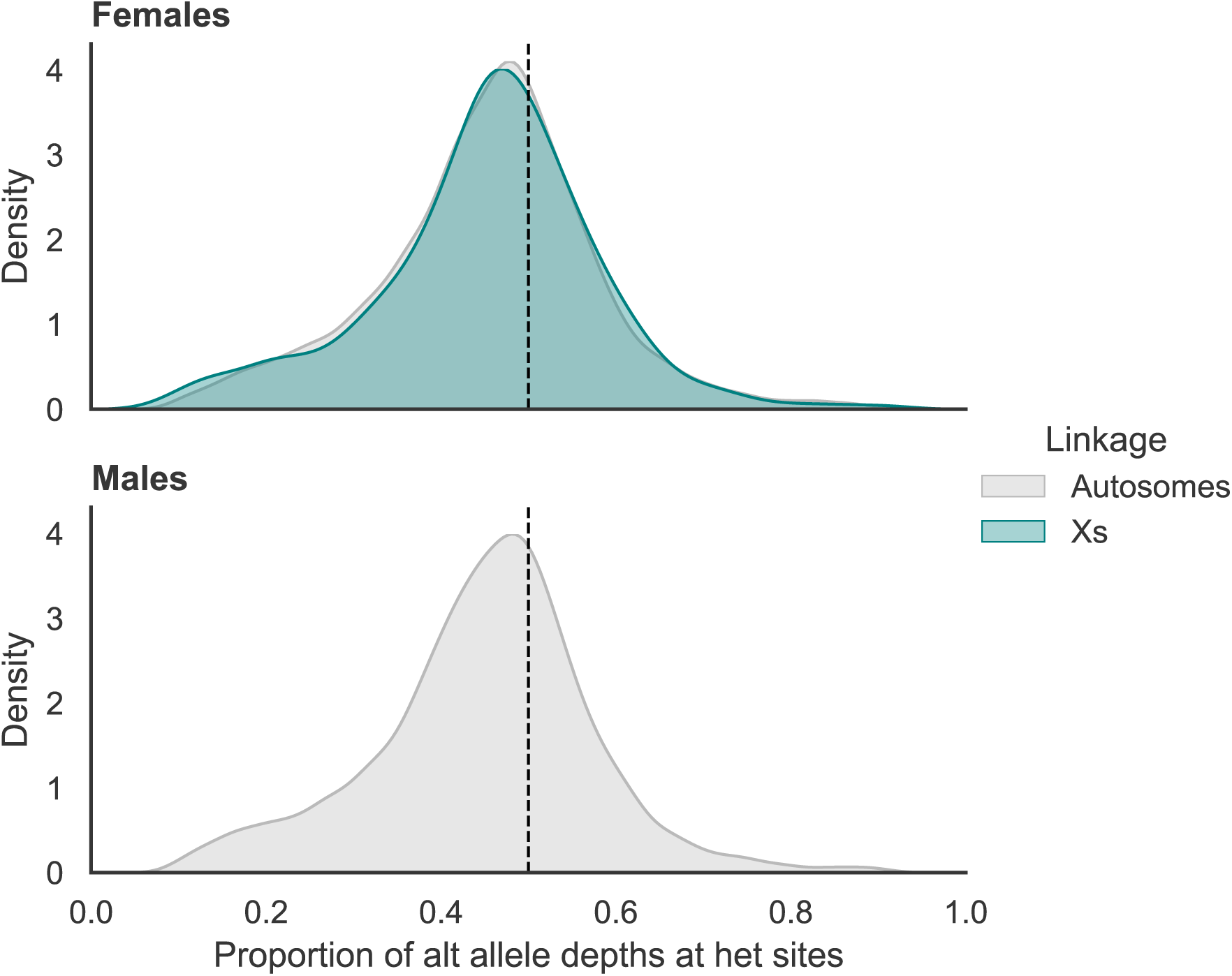
Intermediate coverage depth fractions of reference and alternative alleles at heterozygous autosomal (grey) and X-linked (teal) sites in female (upper) *Allacma fusca* RNA-Seq datasets are equivalent to male (lower) autosomal sites suggesting that males express both their maternal and paternal alleles despite PGE. The slight bias in coverage away from 50:50 (dotted line) towards the reference allele suggests reference bias.

### No evidence for a DNA methylation system in *A. fusca*

Given that DNA methylation (a type of epigenetic mechanism) has been implicated in PGE in an insect species (*Planococcus citri*, see Bain *et al*. (2021)), and potential differences in epigenetic regulation of the Xs and autosomes could affect their evolutionary rates (Ashe *et al*., 2021), we examined the presence of DNA methylation in *A. fusca* via whole genome EM-Seq. Whilst a CpG o/e analysis revealed the potential for genome-wide low levels of DNA methylation, EM-Seq revealed that *A. fusca* exhibits a loss of DNA methylation in both males and females (Supplementary information 1). *A. fusca* also appears to be lacking the genes required for a DNA methylation system; DNMT3 (de novo DNA methylation) and DNMT1 (DNA methylation maintenance) (Bewick *et al*., 2017). Whilst one gene (g20528) showed similarity with 58 insect DNMT1 genes, upon inspection via InterProScan it appears more similar to DNMT2 (containing only a DNA-methylase domain and lacking all other DNMT1 domains, (Engelhardt *et al*., 2022)). DNMT2 is a tRNA methyltransferase and is not involved in the methylation of DNA. Finally, g19426 showed similarity to 5 insect DNMT3 genes and contains a PWWP domain but lacks the ADD domain usually present in DNMT3 enzymes, consistent with the broader loss of the DNA methylation machinery in *A. fusca*.

### X-linked genetic diversity and divergence is surprisingly low under PGE

Given the expected effective population size and demographic history of autosomes and X chromosomes is identical under PGE we expected estimates of neutral diversity to be approximately equivalent, or perhaps slightly lower on the X chromosomes if increased positive and purifying selection is reducing diversity at neutral sites through linked selective effects. We investigated genetic diversity in *A. fusca* using all genes and divergence between each species using 12,296 single-copy orthologs inferred using Orthofinder (Emms & Kelly, 2019). The genomes are broadly syntenic between the autosomes and X chromosomes suggesting no large interchromosomal rearrangements but numerous inversions and small translocations (Fig. S1). Genetic diversity (*π*) is low on the X chromosomes relative to the autosomes suggesting selection may be more effectively fixing and/or eliminating new X-linked variation (Table 2, Fig. S4). Surprisingly, this is true for both nonsynonymous and synonymous diversity, as synonymous X-linked diversity (*π_s_*) is five times smaller than autosomal, and indeed many X-linked genes showed no variation at all (Table 2). It may be that strong selective sweeps or background selection are reducing putative neutral sites to a strong degree, however a difference of similar magnitude is also seen between X-linked and autosomal intergenic sites where we expect linked selection to be weaker (Fig. S5). Additionally, as the estimates of *Froh* (using runs of homozygosity *>* 100kb) are overall very low on both X chromosomes and autosomes (*≈* 2% and *≈* 1% respectively) it does not seem to be due to inbreeding. As with our observed X chromosome hyperexpression, this low X/A diversity ratio was also observed in sciarid flies (Baird *et al*., 2025), hinting this may be driven by some unknown aspect of PGE. In addition, both *d_n_/d_s_* and *ds* are also low on the X chromosomes relative to the autosomes (Table 3, Fig. 4), and as there was no difference in the frequency of optimal codon usage between these chromosomes (Table 3, Fig. S6), this suggests that low X-linked synonymous divergence is not being driven by stronger constraint on codon usage.

**Figure 4:**
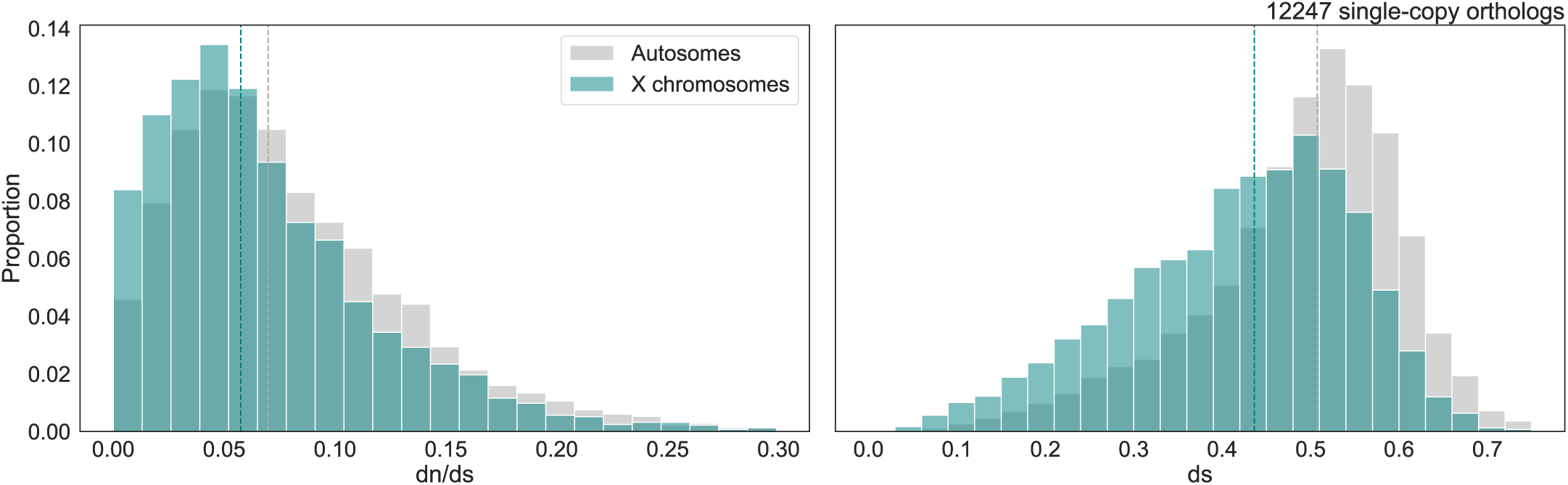
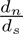 (left) and *d_s_* (right) estimated from 12247 single-copy orthologs across autosomes (grey) and X chromosomes (teal) between *Allacma fusca* and *Sminthurus viridis*.

**Table 2:** Mean pairwise nonsynonymous (*π_n_*) and synonymous (*π_s_*) genetic diversity estimates, and % of genes with zero genetic diversity on autosomes (A) and sex chromosomes (X) in the globular springtail *Allacma fusca*. Values were estimated from 18542 autosomal and 5701 X-linked genes. Confidence intervals are given in square brackets.

| | $\pi_n$ | $\pi_s$ | % $\pi=0$ genes |
| --- | --- | --- | --- |
| **A** | 0.00167 [0.00162, 0.00173] | 0.00556 [0.00541, 0.00570] | 37.3 |
| **X** | 0.000349 [0.000299, 0.000400] | 0.000978 [0.000843, 0.00111] | 73.5 |

**Table 3:** Estimates of mean synonymous (*d_s_*) and nonsynonymous (*d_n_*) genetic divergence on the autosomes (A) and X chromosomes (X) calculated using pairwise alignments of single-copy orthologues (*N_A_* = 8172, *N_X_* = 4075) between the globular springtails *Allacma fusca* and *Sminthurus viridis*. Estimates of adaptive substitution rates are given from averages of per-gene McDonald-Kreitman tests (*α_mckr_*) and from dfe-alpha (Keightley & Eyre-Walker, 2007) (*α_dfe−alpha_*, *ω_α_*). The probability of fixation of deleterious mutations (*P* (*fix_del_*)) is estimated by dfe-alpha. We estimate codon usage bias by the fraction of optimal codon usage (*F_op_*). Confidence intervals are given in square brackets.

| | $d_n$ | $d_s$ | $d_n/d_s$ | $\alpha_{mckr}$ | $\alpha_{dfe-alpha}$ | $\omega_\alpha$ | $P(fix_{del})$ | $F_{op}$ |
| --- | --- | --- | --- | --- | --- | --- | --- | --- |
| **A** | 0.0398 [0.0392, 0.0404] | 0.487 [0.485, 0.490] | 0.0820 [0.0807, 0.0832] | 0.197 [-4.24, 1] | -2.354 [-2.367, -2.348] | -0.108 [-0.109, -0.108] | 0.00131 | 0.429 |
| **X** | 0.0293 [0.0285, 0.0300] | 0.419 [0.415, 0.423] | 0.0692 [0.0675, 0.0708] | 0.868 [0.103, 1] | 0.634 [0.624, 0.644] | 0.0292 [0.0287, 0.0296] | 0.000142 | 0.424 |

### The X chromosome shows both stronger adaptive and purifying selection

We inferred the dDFE (assuming a gamma distribution using a folded SFS) using zero-fold degenerate sites at SCOs as selected sites and intergenic sites in *A. fusca* as putatively neutral sites to infer demography. Given we expect an equivalent demographic history between autosomes and X chromosomes, and weaker background selection on the autosomes, we inferred demography for all chromosomes using only the autosomes (see Methods). Our autosomal intergenic SFS (Fig. S7) was best supported by a three-epoch demographic model (LRT, p *<* 0.05) which indicated a bottleneck (N1/N2 = 10, N1/N3 = 1.43, t2 = 12.5, t3 = 11.7). The inferred dDFE had a smaller shape parameter on the autosomes (*β* = 0.0919) than on the X chromosomes (*β* = 0.260), and X-linked mutations showed a higher proportion of strongly deleterious effects (*NeS >* 100) and a reduced fraction of nearly neutral mutations relative to autosomes (Fig. 5A).

**Figure 5:**
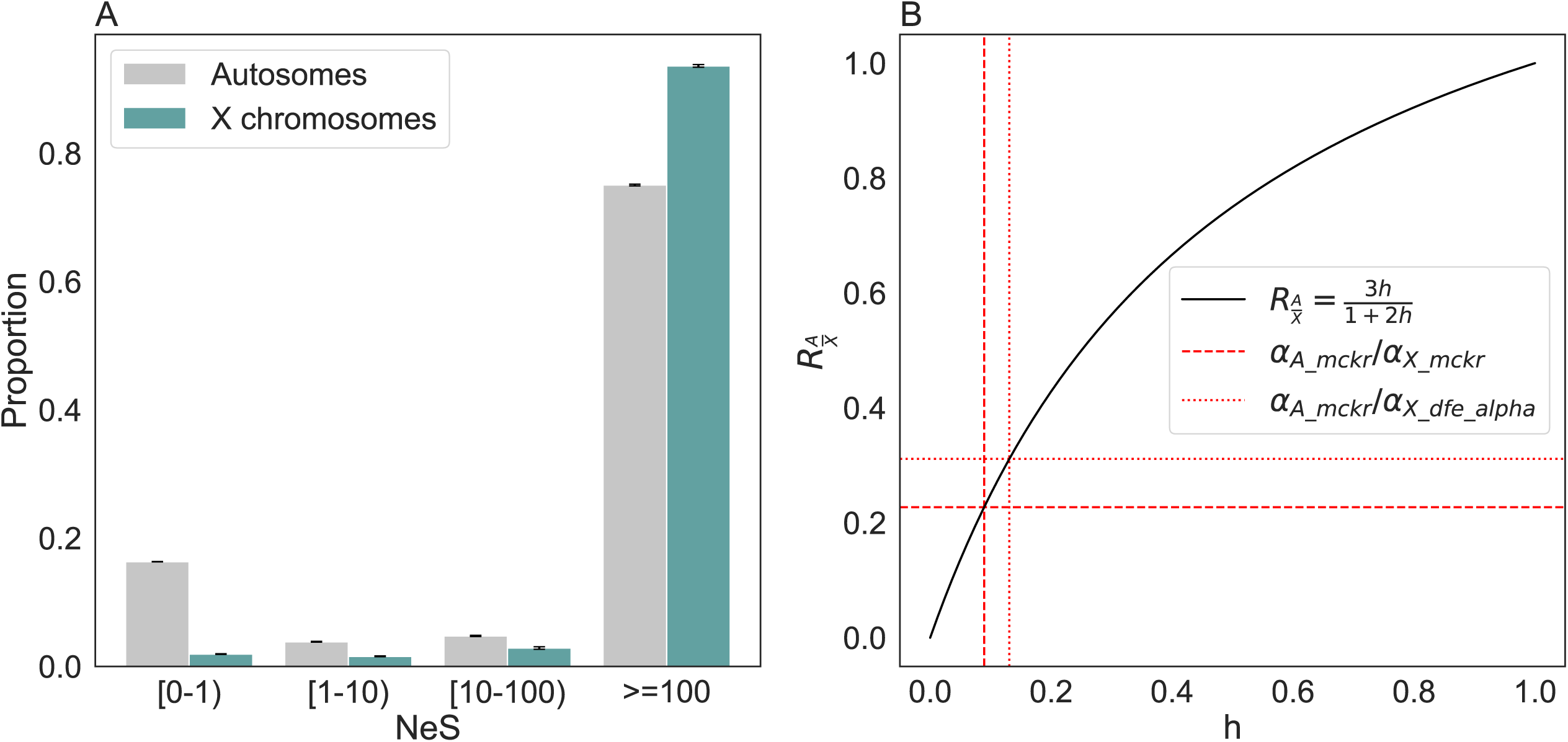
A) The deleterious distribution of fitness effects (dDFE) on the X chromosomes (teal) and autosomes (grey) in *A. fusca* estimated using dfe-alpha. On the x axis, the population-scaled strength of deleterious selection (*N_e_S*) is binned logarithmically with inclusive lower values and exclusive upper values, and increases in strength towards the right. B) Estimates of the mean dominance coefficient (*h*) using ratios of estimates of the strength of positive selection between autosomes and X chromosomes (*R_A/X_*). Point estimates of *h* based on expectations of *R_A/X_*as a function of *h* (assuming PGE and sex-averaged fixation probabilities, see Hitchcock *et al*. (2024)) suggest low average dominance coefficients. Estimates of *R_A/X_* on the autosomes and X chromosomes use either mean values of *α* from the McDonald-Kreitman test or dfe-alpha. Approximate *R̄_A/X_* using estimates of *α_X_* from the McDonald-Kreitman test and dfe-alpha correspond to *h̄ ≈* 0.089 and *≈* 0.13 respectively.

Adaptive substitution rates inferred from both the McDonald-Kreitman test (ns) and DFE-alpha are greater on the X chromosomes than the autosomes (Table 3). Conversely, the probability of fixing a deleterious mutation was considerably higher on the autosomes than the X chromosomes (*P* (*fix_del_*) *A/X ≈* 9.23; Table 3). These both suggest that natural selection is more effective at removing deleterious variation and positively selecting adaptive variation at hemizygous loci due to haploid expression, supporting the faster-X effect.

Given the equivalence of autosome and X chromosome transmission dynamics in PGE systems, we can use these adaptive substitution rates to estimate the mean dominance coefficient (*h*) with much fewer assumptions than typical diplodiploid systems. Assuming new beneficial substitutions occur on the autosomes and X chromosomes at rates *α_A_* and *α_X_* respectively (averaging sex-limited fixation probabilities): *α_A_ ≈* 3*Nµhs* and *α_X_ ≈ Nµs*(1 + 2*h*), hence we can solve 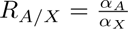 for *h* (Charlesworth *et al*., 1987, Hitchcock *et al*., 2024). Using estimates from the McDonald-Kreitman test and DFE-alpha respectively we estimate low mean dominance coefficients of approximately 0.089 and 0.13 (Fig. 5B).

As the faster-X effect must act on genes expressed by males, we expect it to be more effective at genes where it is expressed mostly by males and less effective at predominately female genes. To test this, we compared *dn/ds* and *α* using the McDonald-Kreitman test at unbiased, male-biased, and female-biased genes between the autosomes and sex chromosomes. Both *dn/ds* (Mann-Whitney U; UB: 0.0685, MB: 0.0915, p *<* 0.001) and *α* (Mann-Whitney U; UB: -0.771, MB: -0.0809, p *<* 0.001) are higher in male-biased genes than unbiased genes genome-wide. When comparing sex chromosome and autosomes, we do not see stronger selection in X-linked male-biased genes. Across both unbiased and male-biased genes we see higher *dn/ds* values on autosomes (Mann-Whitney U; UB: 0.07 vs 0.056, p *<* 0.001; MB: 0.093 vs 0.081, p = 0.00609) and higher *α* values on X chromosomes (Mann-Whitney U; UB: -0.88 vs 0.0013, p *<* 0.001; MB: -0.16 vs 0.35, p *<* 0.001). Female-biased genes show higher *α* values on the X chromosomes only (*dn/ds*: 0.081 vs 0.071, p = 0.119; *α*: 0.077 vs 0.48, p = 0.0403). Thus, it does not appear that male-biased genes experience a stronger faster-X effect than other genes. Interestingly, the male X-linked *dn/ds* distribution does hint at bimodality, with the genes underlying the higher value peak also demonstrating high *α* values (*≈* 0.15). This suggests that a subset of male-biased genes could either experience a stronger effect of X linkage (Fig. 2B) or it may simply be that they are more strongly selected.

## Discussion

We utilised globular springtails, a taxon that undergoes paternal genome elimination (PGE), to investigate how sex-specific hemizygosity alters the efficacy of natural selection. Empirical support for the faster-X effect has historically been difficult to interpret because the effect of a haploidy is entangled with other confounding factors, and as a result numerous interpretations have been given for why X chromosomes evolve rapidly (Charlesworth *et al*., 2018, Mank *et al*., 2010b). This heterogeneity is not limited to XY systems: on the Z chromosome, birds show a faster-Z effect attributed primarily to drift from a reduced effective population size (Mank *et al*., 2010a), whereas in silkmoths and monarch butterflies it is driven by more efficient positive selection (Mongue *et al*., 2022, Sackton *et al*., 2014). Systems undergoing PGE offer a unique opportunity to compare X chromosomes and autosomes with fewer confounding differences than in taxa with diplodiploid XY/ZW systems, as transmission dynamics are equivalent between chromosome types. Thus, leveraging this combination of genome-wide haplodiploid inheritance with diplodiploid autosomal expression, we quantified the differences in positive and purifying selection driving autosome and sex chromosome evolution. Contrary to expectations that hemizygosity should lead to higher genetic divergence rates at non-neutral X-linked sites (see Meisel & Connallon (2013)), we saw reduced X-linked divergence. After partitioning the selected components of divergence, we found adaptive substitution rates were higher on the X chromosome, X-linked sites exhibited surprisingly low genetic diversity, and the distribution of deleterious fitness effects was strongly skewed, which signals more efficient purifying selection. Our finding that both purifying and positive selection are more efficient on the X, despite the absence of the demographic differences often invoked to explain faster-X patterns (Mank *et al*., 2010b) solidifies the expression of recessive alleles in haploid males as the driving factor. This converges with results from an independent PGE lineage, Sciarid flies (Baird *et al*., 2025), which likewise found evidence for the faster-X effect. Together, these results suggest that ploidy-driven faster-X evolution may be a more general process than the mixed empirical record from conventional diplodiploid XY/ZW systems has suggested, and that PGE taxa may be an underused resource for testing sex chromosome evolution theory more broadly.

We demonstrated that male globular springtails express both parental alleles on the autosomes, which confirms that the paternal autosomes are transcribed despite their elimination during spermatogenesis. This corroborates cytogenetic observations that paternal autosomes are only eliminated from male germline cells during male meiosis but are retained in all other tissues in a euchromatic state (Herbette & Ross, 2023). Dosage is approximately balanced on all chromosomes between the sexes suggesting that there is full dosage compensation of the X through the upregulation of the X in males, similar to mechanisms found in *Drosophila* for example (Disteche, 2012). While we did detect a slight skew between sexes due to very highly expressed male-biased genes, there was evidence of modest X hyperexpression in both males and females. This phenomenon has been reported in another PGE system (Baird *et al*., 2025) and in some diplodiploid species as a dosage compensation mechanism (Allen *et al*., 2013, Prince *et al*., 2010). It may be that overcompensation occurs due to imperfect upregulation and selection favouring increased expression in both sexes (Charlesworth & Charlesworth, 2025, Prince *et al*., 2010). Regardless, these expression patterns show that reduced X-linked expression due to hemizygosity is not a constraint on faster-X evolution in this system. Given the expression distributions on the autosomes and sex chromosomes are very similar between males and females, dosage compensation may occur through a chromosome-specific rather than copy-number specific transcription property (Charlesworth & Charlesworth, 2025). Toups & Vicoso (2023) suggest that one of the symphypleonan X chromosomes may share a common ancestor with the ancestral Insecta X, but the origin of the second X chromosome is currently unknown, and to date no Y chromosomes have been observed. Our finding that dosage is more unbalanced on the second X chromosome does hint that it may be younger. In hexapods, DNA methylation is primarily located in gene-bodies and is associated with stable gene transcription (Dixon & Matz, 2022). However, breaking this trend, DNA methylation has been suggested to be involved in the silencing of the paternal chromosomes in males of the mealybug, *P. citri* (Bain *et al*., 2021). As this is currently the only Arthropod to exhibit this particular type of DNA methylation profile, we hypothesized that sex-specific DNA methylation may be a conserved feature of species with PGE. Whilst we find that *A. fusca* lacks a DNA methylation system, this is somewhat consistent with our finding that paternal genes are expressed, since NA methylation may be involved in paternal gene silencing in mealybugs. More broadly, our study is the first to examine the DNA methylation profile of a species of springtail and documents the first confirmed loss in Collembola, although previous work in *Holacanthella duospinosa* is suggestive of a lack of DNA methylation (Wu *et al*., 2017).

Concordant with our results, it has frequently been observed that male-biased genes evolve most rapidly (Ellegren & Parsch, 2007), and it may be that this is due to the accumulation of male-biased genes on sex chromosomes in response to sexual antagonism (Ellegren & Parsch, 2007), in concert with the faster-X effect (Baines *et al*., 2008). While we see a substantial excess of male-biased compared to female-biased genes, including an accumulation on the X chromsomes, we see no evidence of more effective natural selection due to X-linkage in any class of sex-bias, suggesting that faster-X evolution is likely to be principally underpinned by a different or narrower subset of genes. This corroborates previous evidence from a PGE system which also found slower X-linked *dn/ds* across all classes of sex-bias (Baird *et al*., 2025). Instead, a higher genome-wide *α* suggests that male-biased genes on average experience more positive selection as a whole. We note that as our data are restricted to whole-adults, and proportions of germ tissue vary between sexes, the bias and selection profiles observed here may differ from younger stages of life history.

We see reduced genetic diversity and divergence on the X chromosomes. As both selected and neutral X-linked genetic diversity is particularly low, *d_N_ /d_S_* ratios are reduced, and the inferred distribution of deleterious fitness effects is skewed (assuming X-linked and autosomal genes have equivalent fitness effects in new mutations), purifying selection is likely more efficient on the X chromosomes as predicted due to hemizygosity. Reduced *d_N_ /d_S_* rates might be surprising given we expect an increased adaptive substitution rate on the X chromosomes. However, we also expect that the weakly deleterious substitution rates should be higher on the autosomes, and given that the majority of new mutations are deleterious (Charlesworth & Charlesworth, 1998), it seems likely that the net effect is a relatively higher autosomal substitution rate. In contrast to typical diplodiploid systems, reduced X-linked diversity in globular springtails is not a consequence of differences in effective population size, as the autosomes and X chromosomes share an identical inheritance under PGE. Instead, low X-linked diversity is most likely a result of selection or mutation-rate differences. Reduced neutral diversity and divergence may reflect more effective X-linked background selection and more frequent selective sweeps. It may even be that the X-linked gene set is more conserved and thus experiences stronger background selection due to function and not simply ploidy, but without stronger constraint on codon usage (Table 3, Fig. S6). Low synonymous divergence on the X suggests a lower mutation rate may also have contributed to low divergence and diversity. This may emerge from selection favouring mutation rate modifiers on hemizygous chromosomes to reduce new mutation load in males (Charlesworth & Charlesworth, 1998, McVean & Hurst, 1997). Even if the difference in synonymous divergence rates between chromosomes entirely reflects mutation rate differences in absence of background selection, the magnitude of this difference (*X/A ≈* 0.9) is much smaller than the difference in genetic diversity (*X/A ≈* 0.2), and local diversity is accounted for when inferring adaptive substitution, so it cannot exclusively account for these observed differences. Lastly, low X chromosome genetic diversity and neutral divergence does not appear to be a consequence of inbreeding in *Allacma fusca* given no significant runs of homozygosity.

The expected mean dominance coefficient (*h*) can be derived as a function of the ratio of adaptive substitution rates between autosomes and X chromosomes (*R_A/X_*) (Charlesworth *et al*., 1987, Hitchcock *et al*., 2024). It is particularly appealing in globular springtails because recombination rates, effective population sizes, demographic histories, and the relative importance of selection between sexes are equivalent between all chromosomes under PGE, and thus interpretation is grounded on fewer assumptions. Using estimates of the adaptive substitution rate from the McDonald-Kreitman test and DFE-alpha, we inferred *h̄* as 0.089 and 0.13 respectively, consistent with predominantly recessive beneficial alleles (Fig. 5B). Although in PGE systems the faster-X effect is expected under all values of *h*, these results suggest that new mutations are mostly recessive, and faster-X effects are indeed driven by hemizygosity enhancing the efficacy of both purifying and positive selection. However, if genetic diversity is strongly reduced due to haplodiploid transmission, as a consequence of increased background selection, sweeps, and selection for reduced X-linked mutation rates, the observable impact of faster-X adaptation in PGE systems could be limited.

## Conclusion

The dynamics of paternal genome elimination provide powerful natural experiment for studying the principles of population genetics given they show a distinct set of characters compared to diplodiploid system. We demonstrate this by quantifying the faster-X effect and mean dominance coefficient in globular springtails. This estimate was based on the assumption that the large number of genes on X chromosomes and autosomes are comparable with respect to strength of selection and dominance, which might not be fully met, and so further advances would require estimating within homologous genes and preferentially with numerous biological replicates. Additionally, while sharing a single demographic model between X chromosomes and autosomes removes one of the principal confounding factors, our inference of the deleterious DFE itself was still performed under an assumption of additivity. Given that our results suggest new mutations deviate substantially from additivity, methods that jointly co-infer the DFE and h remain strongly desirable, and PGE systems may offer particularly clean systems for applying them.

We discovered a dramatic reduction of genetic diversity on the X chromosmes of *A. fusca*, which is already the second PGE system with this pattern. We currently lack a good explanation for the phenomenon and this opens up the question of whether it is the consequence of dramatically different mutation rates between chromosomal sets, or some other principle associated with PGE. Our results support the faster-X effect as hemizygosity facilitates both purifying and positive selection, but the former dominates as weak but frequent deleterious substitutions are more likely to accumulate on the autosomes due to genetic drift, leading to a net reduction in X-linked divergence. Comparative work across independent instances of PGE will be key to understanding why PGE evolves in the first place, how it affects chromosome evolution, and whether the patterns we see here are a general consequence of PGE or something more specific to this lineage. These systems may also shed light on the role of sex chromosomes in speciation — particularly Haldane’s rule and the large-X effect. More broadly, direct estimates of the dominance coefficient of new mutations from natural populations remain rare, having so far been obtained in only a handful of taxa using methods that are often confounded by demography or mating system. By exploiting the equalised demographic and recombination context that PGE provides, our estimate of *h̄* offers a comparatively assumption-light estimate conditioned on the same background DFE, and adds a rare data point to a still poorly resolved area of population genetics.

## Methods

### Sampling and sequencing data

We used the *A. fusca* reference genome (GCA 947179485.1) for the focal species analysis (Jaron *et al*., 2023) and the reference genome of *Sminthurus viridis*, a relative within the Sminthuridae (GCA 965194885.1) as an outgroup, both generated within the Darwin Tree of Life project (of Life Project Consortium, 2022). Sequence evolution analyses were conducted using our previously generated resequencing data of 12 individuals of *A. fusca* sampled in Wytham woods, Oxford and Holyrood park, Edinburgh Jaron *et al*. (2022).

To annotate the genome, analyse biparental, sex-biased expression and dosage compensation, we sequenced 20 RNA-Seq datasets for the globular springtail *Allacma fusca* (see Table S1). Sampling was conducted in the field from 2018 to 2021 in Scotland, England, and The Netherlands. Captured individuals were kept individually in plaster cages and daily screened for presence of spermatophores to identify males. Females were identified as individuals without generating spermatophores for three consecutive days and using presence of subannal appendage and larger body size as an additional evidence. The sex of all individuals was confirmed genomically. The RNA-Seq data consisted of ten female and ten male datasets, five of each were generated from pools of three individuals due to input limitations (of samples collected in 2018), while the later generated RNA-Seq was generated from single individuals. All samples were collected from surfaces using aspirators and were frozen live in liquid nitrogen. RNA-extractions from pools was performed using the Purelink RNA Purification kit protocol, and TruSeq stranded mRNA-Seq libraries were generated with poly-A selection and sequenced on an Illumina NovaSeq platform. The five individual male and female *A. fusca* samples were processed using dual RNA and DNA extraction using the Qiagen AllPrep DNA/RNA Mini Kit. Individuals were crushed using a manual eppendorf pestle in Buffer RTL before continuing with the manufacturer’s protocol for animal tissues. Resulting DNA and RNA were treated with RNase A and DNase I respectively. The DNA part of the extraction was used for EM-Seq, while the RNA libraries were constructed using TruSeq stranded mRNA-Seq library. EM-Seq and RNA-Seq library preparation were carried out by the NERC Environmental Omics Facility and libraries were sequenced on an Illumina HiSeq 2500 platform respectively using 150bp paired-end reads.

### QC and genome annotation

WGS reads were trimmed using skewer v0.2.2 (Jiang *et al*., 2014) and RNA-Seq reads were trimmed using FastP (Chen *et al*., 2018). Read quality was assessed for both datasets using FastQC (Andrews *et al*., 2010). Each RNA-Seq dataset was mapped individually to the *A. fusca* reference genome using HISAT2 (Kim *et al*., 2019) for gene annotation.

Repetitive elements were annotated in each reference genome using EarlGrey (Baril *et al*., 2024). Following this, genes were annotated using BRAKER (Gabriel *et al*., 2024) on each repeat-softmasked genome. For *A. fusca* we ran the BRAKER3 pipeline using the RNA-Seq data as hints and the OrthoDB v.11 arthropoda partition generated for use with BRAKER as a reference protein database (Kuznetsov *et al*., 2023). For *S. viridis* we ran the BRAKER2 pipeline using the aforementioned OrthoDB database combined with the protein sequences predicted in *A. fusca*. Both gene annotation files generated by braker were processed using AGAT (Dainat, 2022).

### Variant calling

Trimmed WGS reads were mapped to the *A. fusca* genome using bwa mem (Li, 2013). Duplicates were marked in each read group using Sambamba (Tarasov *et al*., 2015). We used mosdepth (Pedersen & Quinlan, 2018) to annotate callable regions based on a minimum coverage of 8 and a maximum coverage of twice the mean of the coverage distribution. We intersected regions callable in all datasets and removed repetitive regions used BEDTOOLS (Quinlan & Hall, 2010), to identify regions suitable for variant calling. We called variants using Freebayes (Garrison & Marth, 2012) using an expected theta of 0.01, a max complex gap of 1, using mapping quality for data likelihoods, and without Hardy-Weinberg Equilibrium priors. Variants were filtered using BCFtools (Li, 2011) to normalise, decompose allelic primitives, filter for a minimum quality of 10, a minimum SNP gap of 2, and for balance satisfying *^′^RPL >* 1*|RPR >* 1*|SAF >* 1*|SAR >* 1*^′^*. These steps are implemented in a Nextflow pipeline available at github.com/samebdon/var_call.

### Gene Expression

We used RSEM (Li & Dewey, 2011) to quantify gene expression from each RNA-Seq dataset where both reads map to the features described in the gene annotation. We then tested genes for sex-biased gene expression (SBGE) using the DESeq2 package available in R (Love *et al*., 2014). The first principal component of a PCA of the expression data suggests that one of the multi-sample datasets (AF F 5) likely contains a mixture of two males and one female, and was thus labeled as mixed. All remaining samples were correctly identified as males or females morphologically. Genes were labelled as sex-biased if the absolute log2 fold change between males and females was greater than one and supported by an adjusted p-value *<* 0.05.

We investigated whether male globular springtails express both parental haplotypes as expected given their mode of PGE by calling variants with the single-sample RNA-Seq datasets following the aforementioned variant calling pipeline but using Hisat2 (Kim *et al*., 2019) to align mono-sample RNA-Seq reads to the reference. We then quantified coverage support of each allele at heterozygous SNPs (0.1 =*< Ref AD* =*<* 0.9) to demonstrate biparental expression.

We tested for dosage compensation as follows. We mapped the trimmed RNA-Seq reads to transcripts extracted from the reference assembly using RSEM. We normalised expression levels as TMM transcripts per million mapped reads (TMM-TPM) and averaged across males and females per gene. We excluded genes with a mean (per sex) log2(TMM-TPM) below the 2.5% and above the top 97.5% quantiles. We test for dosage compensation by testing whether sex chromosomes and autosomes differ in expression in each sex, and dosage balance by comparing males and females on sex chromosomes and autosomes separately. We further used TMM-TPMs to infer the optimal codons and estimate optimal codon frequency of A-linked and X-linked genes using the R package ‘cubar’. Additionally, we quantified codon usage bias to check whether the faster-X effect could be constraining X-linked divergence at synonymous sites by estimating the frequency of optimal codon usage (Fop) using TMM-TPMs (Campos *et al*., 2013).

### DNA methylation

A CpG observed/expected analysis was carried out on all genes, exons and introns, following: https://github.com/MooHoll/cpg_observed_expected. An observation of less CpGs than expected, given the GC content, is indicative of DNA methylation presence (Bewick *et al*., 2017). EM-Seq data were quality checked using FastQC (Andrews *et al*., 2010) and aligned to the reference genome using Bismark with standard parameters (Krueger & Andrews, 2011). Data were also aligned to the lambda phage genome and pUC19 plasmid genome, which serve as an unmethylated and methylated control respectively, to determine the efficiency of the enzymatic conversion. Methylated calls generated by Bismark did not exceed the conversion error rates for the lambda control (0.1% methylation) and pUC19 control (98.1% methylation) suggesting a lack of DNA methylation in *A. fusca*. To determine the presence of genes required for DNA methylation establishment and maintenance, a reciprocal protein blast was carried out using 321 insect DNMT1 protein sequences and 110 DNMT3 protein sequences from https://v2.insect-genome.com/Pcg, using blastp (Camacho *et al*., 2009) with a minimum e-value of 1e-3. Conserved domains of top hits were identified using InterProScan (Jones *et al*., 2014).

### Estimating the distribution of fitness effects and adaptive substitution rates

We identified single-copy orthologs (SCOs) between our ingroup and outgroup by clustering protein sequences of the longest isoforms of complete proteins with Orthofinder (Emms & Kelly, 2019). To estimate nonsynonymous and synonymous genetic diversity and divergence between SCOs we used a custom script (https://github.com/samebdon/orthodiver) with randomly phased haplotypes aligned using TranslatorX (Abascal *et al*., 2010) and guided by protein alignments for each gene generated with MAFFT (Katoh *et al*., 2002). We generated folded site-frequency spectra (SFS) for nonsynonymous and synonymous sites within SCOs using scikit-allel (Miles *et al*., 2021) after annotating per-site degeneracies with degenotate (Mirchandani *et al*., 2024). We use folded SFSs given a lack of suitable outgroup for inferring ancestral allele states. SFSs were generated for intergenic sites and SCOs on the autosomes and on the X chromosomes respectively.

We used DFE-alpha (Eyre-Walker & Keightley, 2009, Keightley & Eyre-Walker, 2007) to estimate the difference in adaptive substitution rates between the autosomes and X chromosomes while accounting for the deleterious distribution of fitness effects (DFE). For estimating the DFE and alpha on both the autosomes and the X chromosomes we fit a three epoch demographic model to the intergenic autosomal SFS for three reasons. Firstly given that we expect the same demographic history in each chromosome class given paternal genome elimination we can base the inference for each on the same SFS. Secondly, as DFE-alpha assumes sites are neutral and unlinked to infer demographic history we use intergenic sites to minimise linked selective effects (using synonymous genic sites to infer demography can lead to the false inference of population expansions (Messer & Petrov, 2013)). Lastly, we exclude X-linked intergenic sites as we expect these sites will violate the assumption of neutrality to a greater degree than autosomal sites given the expected consequences of hemizygosity. We then inferred the deleterious DFE (assuming a gamma distribution) for the autosomes and X chromosomes respectively using zerofold-degenerate sites in single-copy orthologs given the shared inferred demographic history. Adaptive substitution proportions (*α*) and relative rates (*ω_α_*) were then estimated for the autosomes and X chromosomes separately given their respective deleterious DFEs and divergence estimates from fourfold degenerate sites whilst applying a Jukes-Cantor correction and removing polymorphism contributing to divergence. Intergenic sites were not used to estimate divergence for this stage of the inference as divergence between the ingroup and the outgroup is too great to generate reasonable alignments. This was initially run for all genes and then unbiased and male-biased genes separately.

## Acknowledgements

This research was funded in whole, or in part, by the Wellcome Trust 220540/Z/20/A. For the purpose of Open Access, the author has applied a CC BY public copyright license to any Author Accepted Manuscript version arising from this submission.

The single-individual RNA-Seq and EM-Seq was funded by the UK Natural Environment Research Council (NERC) Environmental Omics Facility, project NEOF1345 awarded to HM and KSJ.

## Data Availability Statement

Raw RNA-Seq reads are deposited at the ENA under accession number PRJEB126531. All EM-Seq data are available at NCBI, under BioProject number PRJNA1461370. All WGS data is from ENA/SRA under accession PRJEB44694.

