## Supplementary materials for "Stronger positive and purifying selection on X-linked genes in a species with paternal genome elimination"

### Supplementary information 1

The CpG o/e profile of A. fusca (Fig. S8) shows a unimodal distribution with a skew for a lower CpG o/e in introns. Generally, in Arthropods a unimodal distribution is suggestive of a lack of DNA methylation ((Bewick *et al.*, 2017)), however this does not hold true for all species. For example, the cockroach Blatella germanica, exhibits a similar CpG o/e profile ((Thomas *et al.*, 2020)) and has high levels of genome-wide DNA methylation ((Lewis *et al.*, 2020)). However, EM-Seq of individual males and females reveals DNA methylation levels within the bounds of the errors rates (Table S1). The average enzymatic conversion rate was 0.1% false positives (lambda) and 1.9% false negatives (pUC19). Whilst the typical EM-Seq false negative rate could obscure low levels of DNA methylation in A. fusca, the lack of DNMT genes is suggestive of a complete loss.

24

### Supplementary tables and figures

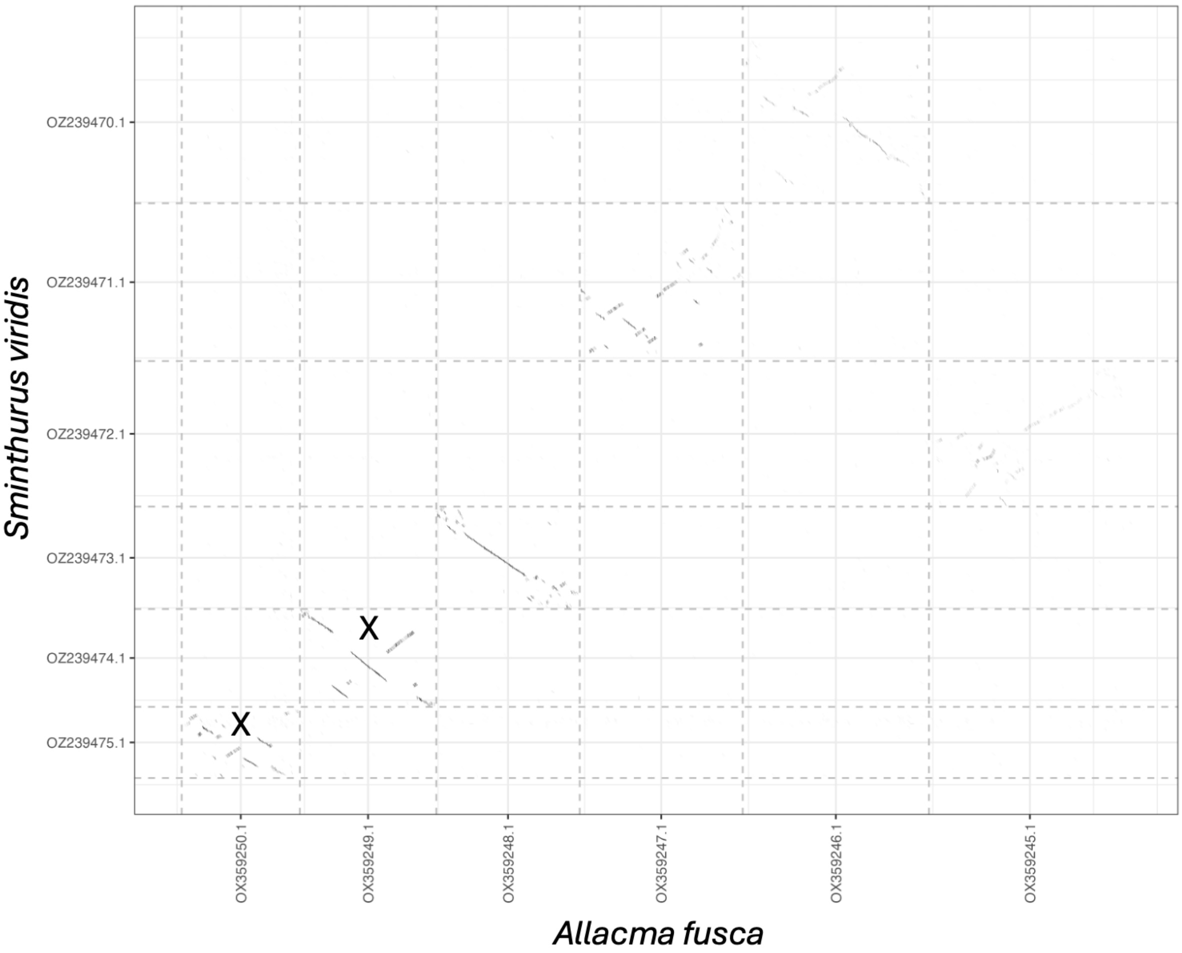


Figure S1: Dot plot between *Allacma fusca* and *Sminthurus viridis*. X chromosomes are annotated on the plot. All other chromosomes are autosomes.

Table S1: DNA methylation levels of the lambda spike, pUC19 spike and per sample methylation levels, shown as the percentage of cytosines found to be methylated.

| **Sample** | **Sex** | **Lambda CpG Methylation (%)** | **pUC19 CpG Methylation (%)** | **CpG Methylation (%)** | **CHG Methylation (%)** | **CHH Methylation (%)** |
|---|---|---|---|---|---|---|
| AF50 | Male | 0.2 | 98.4 | 0.1 | 0.1 | 0.1 |
| AF51 | Male | 0.1 | 97.8 | 0.2 | 0.2 | 0.2 |
| AF53 | Male | 0.2 | 97.8 | 0.2 | 0.2 | 0.2 |
| AF54 | Male | 0.2 | 98.1 | 0.1 | 0.1 | 0.1 |
| AF55 | Male | 0.1 | 98.1 | 0.1 | 0.1 | 0.1 |
| AF58 | Female | 0.1 | 98.3 | 0.1 | 0.1 | 0.1 |
| AF64 | Female | 0.2 | 98.6 | 0.2 | 0.1 | 0.2 |
| AF65 | Female | 0 | 98 | 0.2 | 0.1 | 0.1 |
| AF72 | Female | 0.2 | 97.7 | 0.2 | 0.2 | 0.2 |
| AF74 | Female | 0.1 | 98.3 | 0.2 | 0.2 | 0.2 |

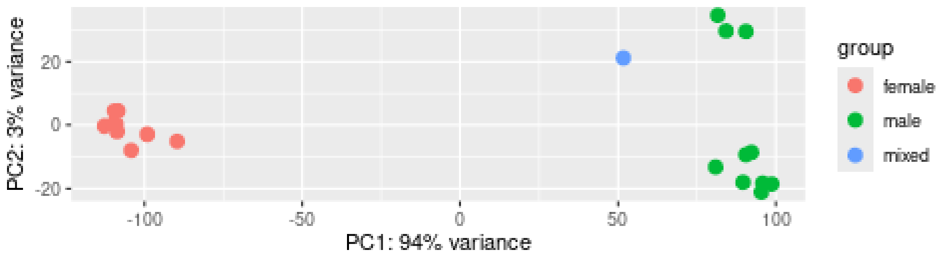


Figure S2: PCA of *Allacma fusca* RNAseq datasets generated in DESeq2.

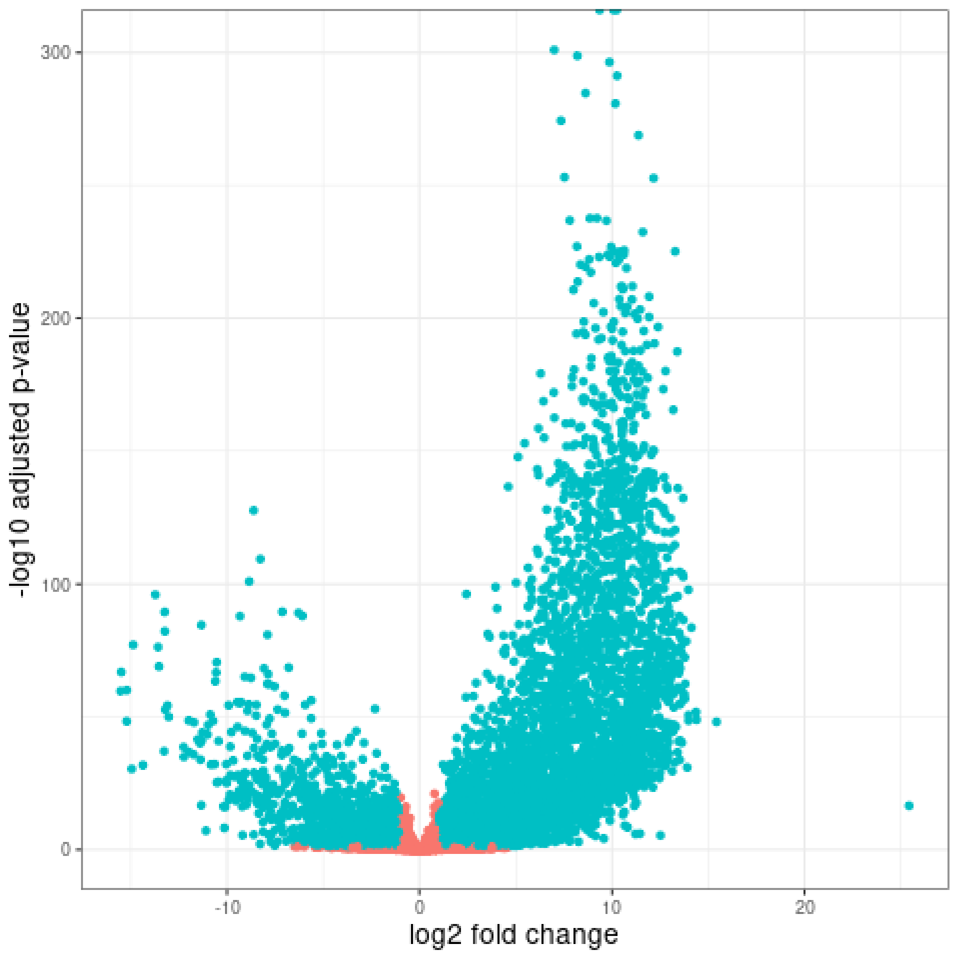


Figure S3: Sex-biased (blue) and unbiased (red) genes in *Allacma fusca* identified using DESeq2. Male biased genes have positive and female biased genes negative log2 fold changes.

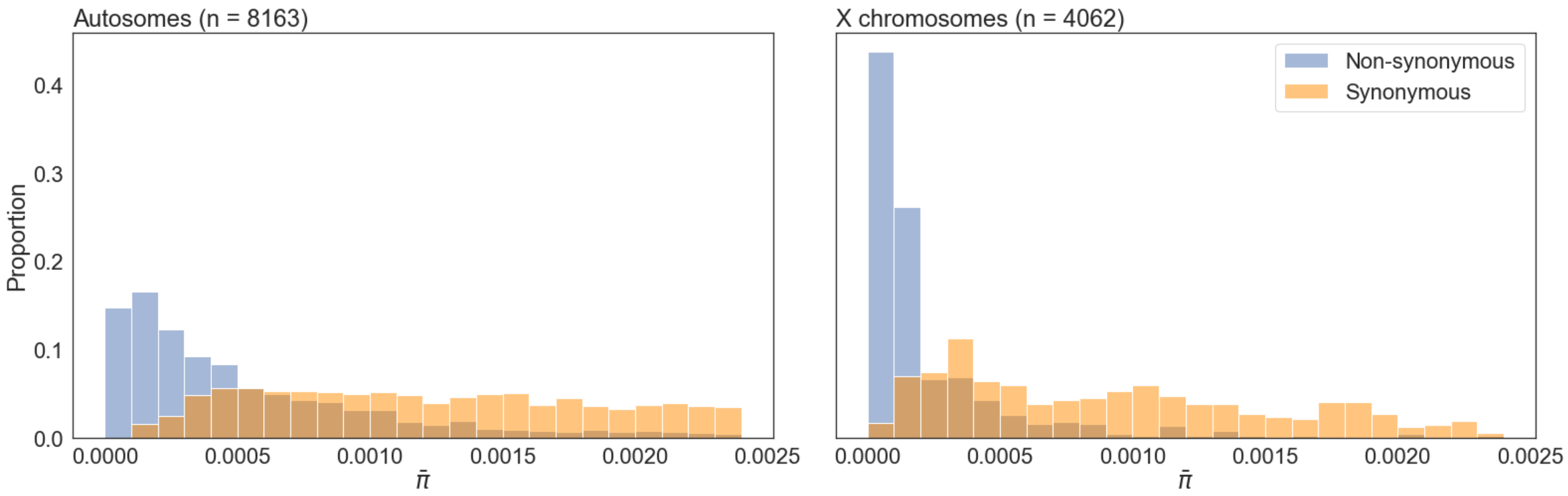


Figure S4: Nonsynonymous and synonymous genetic diversity ($\pi$) in autosomal (left) and X-linked genes (right) in *Allacma fusca*.

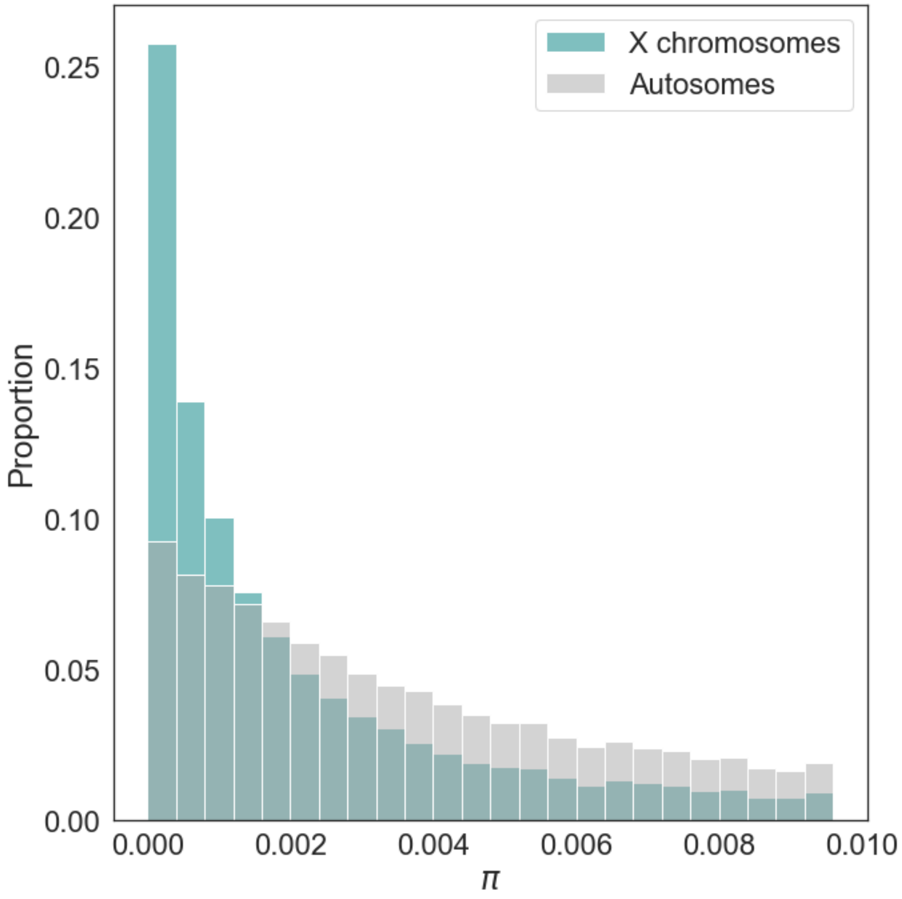


Figure S5: Intergenic genetic diversity ($\pi$) on the autosomes and X chromosomes in *Allacma fusca*.

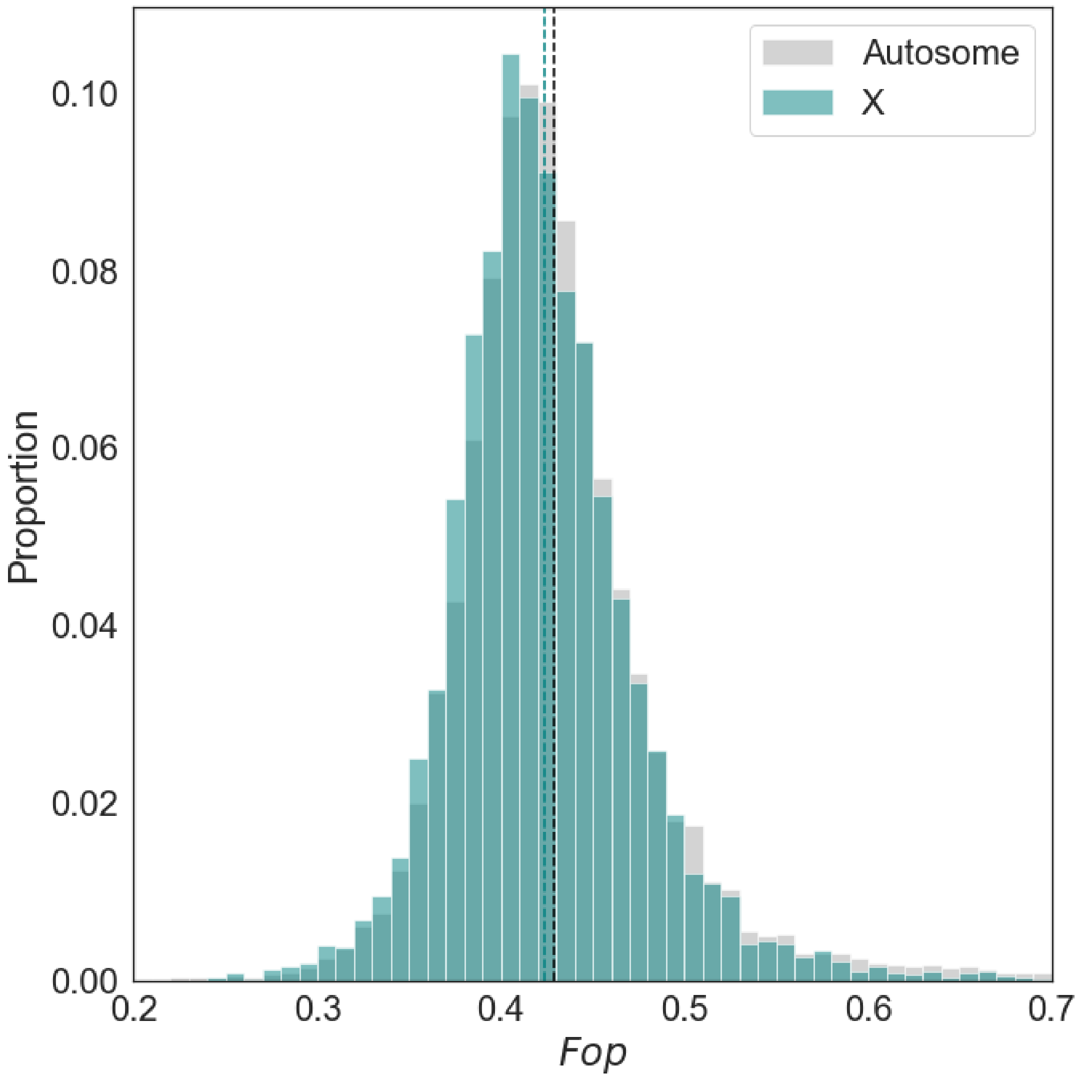


Figure S6: Frequency of optimal codon usage (Fop) on the autosomes and X chromosomes in *Allacma fusca*.

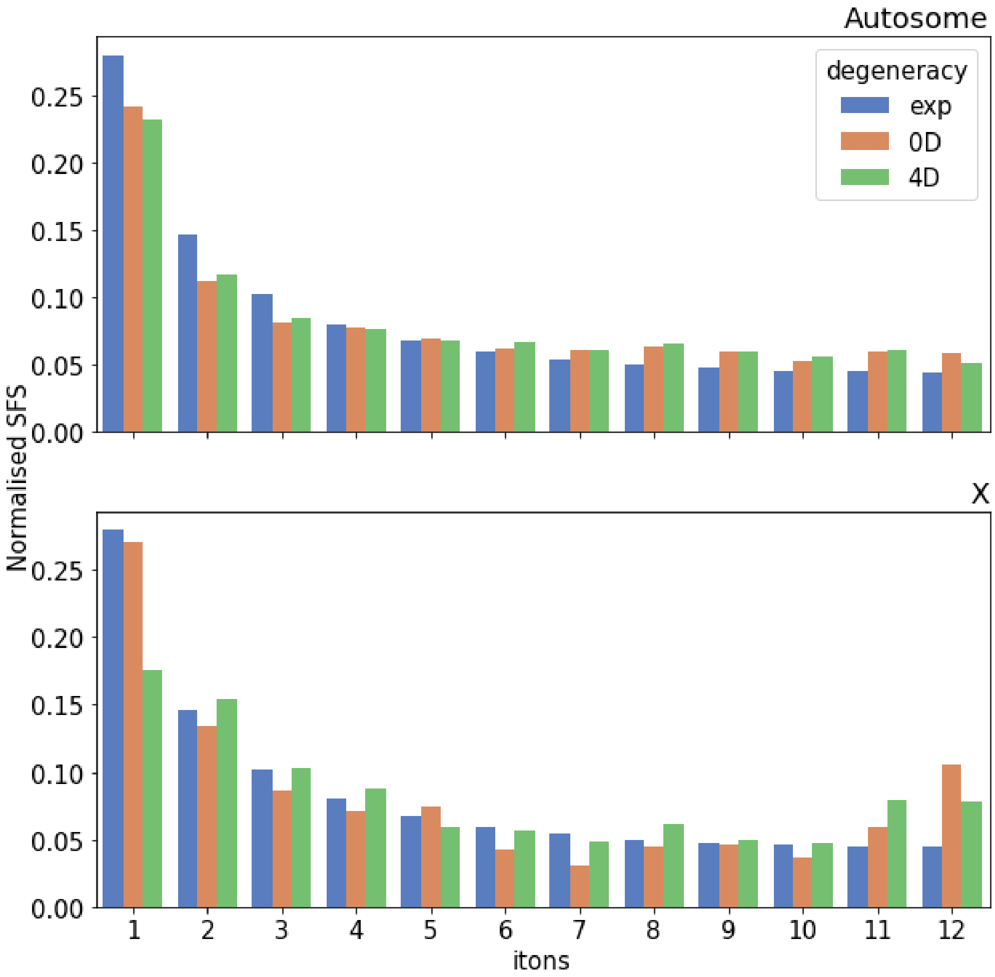


Figure S7: Normalised folded site-frequency spectra at nonsynonymous and synonymous autosomal (upper) and X-linked (lower) sites in *Allacma fusca* relative to a neutral Wright-Fisher expectation.

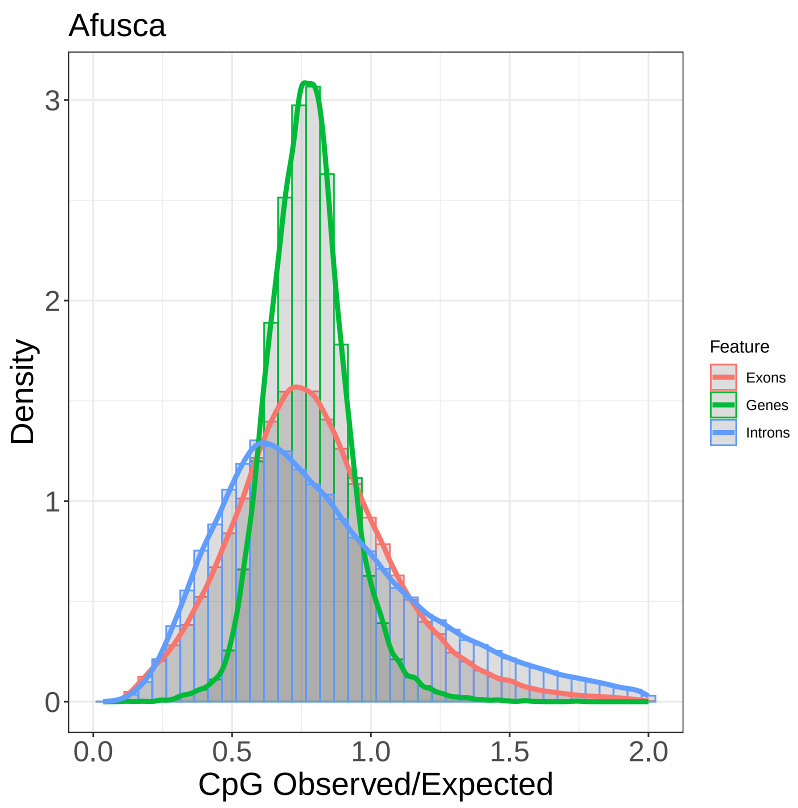


Figure S8: CpG observed/expected distribution for all genes (n = 23,880), exons (n = 139,975) and introns (n = 117,106) in the *A. fusca* reference assembly.
